# SARS-CoV-2 Fusion Peptide has a Greater Membrane Perturbating Effect than SARS-CoV with Highly Specific Dependence on Ca^2+^

**DOI:** 10.1101/2021.01.04.425297

**Authors:** Alex L. Lai, Jack H. Freed

## Abstract

Coronaviruses are a major infectious disease threat, and include the zoonotic-origin human pathogens SARS-CoV-2, SARS-CoV, and MERS-CoV (SARS-2, SARS-1, and MERS). Entry of coronaviruses into host cells is mediated by the spike (S) protein. In our previous ESR studies, the local membrane ordering effect of the fusion peptide (FP) of various viral glycoproteins including the S of SARS-1 and MERS has been consistently observed. We previously determined that the sequence immediately downstream from the S2’ cleavage site is the *bona fide* SARS-1 FP. In this study, we used sequence alignment to identify the SARS-2 FP, and studied its membrane ordering effect. Although there are only three residue difference, SARS-2 FP induces even greater membrane ordering than SARS-1 FP, possibly due to its greater hydrophobicity. This may be a reason that SARS-2 is better able to infect host cells. In addition, the membrane binding enthalpy for SARS-2 is greater. Both the membrane ordering of SARS-2 and SARS-1 FPs are dependent on Ca^2+^, but that of SARS-2 shows a greater response to the presence of Ca^2+^. Both FPs bind two Ca^2+^ ions as does SARS-1 FP, but the two Ca^2+^ binding sites of SARS-2 exhibit greater cooperativity. This Ca^2+^ dependence by the SARS-2 FP is very ion-specific. These results show that Ca^2+^ is an important regulator that interacts with the SARS-2 FP and thus plays a significant role in SARS-2 viral entry. This could lead to therapeutic solutions that either target the FP-calcium interaction or block the Ca^2+^ channel.

## Introduction

Coronaviruses (CoVs) are a major infectious disease threat for humans and animals; examples include SARS-CoV, MERS-CoV, and SARS-CoV-2^1^ (SARS-1, MERS and SARS-2 for simplicity). The pandemic outbreaks of severe acute respiratory syndrome (SARS) in 2003 and Middle East respiratory syndrome (MERS) in 2012 caused 774 and 624 deaths worldwide respectively^1^. SARS-2 is the pathogen of the ongoing COVID-19 pandemic, which has severely harmed our public health, economy and social life. The study of SARS-2 is one of the central agendas in an effort to combat the pandemic.

Glycoproteins located on the viral envelope are required to mediate the viral entry into the host cells and it is a major pathogenicity determinant^2,3^. The viral spike protein (S) is such a CoV glycoprotein. Like SARS-1 and MERS, the S protein of SARS-2 consists of S1 and S2 subunits. The two major steps in viral entry are 1) receptor binding, in which the S1 subunit recognizes a receptor on the host cell membrane, e.g., ACE2, and attaches the virus to the host cell, followed by 2) membrane fusion, in which the S2 subunit mediates the viral envelop membrane and the host membrane fusing together and releasing the virion into the host cell. Membrane fusion is a required stage in viral entry^4^; thus, the blocking of this membrane fusion could be an objective leading to vaccines and therapies to combat COVID-19^5^. The major region of S that interacts with lipid bilayers of the host is called the “fusion peptide” (FP). The FP is critical for membrane fusion, as it inserts into the host lipid bilayer upon activation of the fusion process, perturbing the membrane structure, and initiating the membrane fusion.

Knowledge of the structure and function of the S of SARS-2 is thus important in order to better understand the process of viral transmission, its mechanism and for the development of medical countermeasures (including vaccines, inhibitory peptides, and drugs). The FP is highly conserved across the diverse CoV family, so it serves as a desirable potential target. Although the S protein is categorized as a class I viral fusion protein, it differs in several ways from a typical class I viral fusion protein. Whereas other class I viral fusion proteins such as influenza and HIV glycoproteins are activated via cleavage by host cell proteases at a single site directly adjacent to the fusion peptide, coronaviruses have two distinct cleavage sites (S1/S2 and S2’) that can be activated by a much wider range of proteases, with FP function able to be modulated by changes in the cleavage site position relative to the FP. This gives coronaviruses unique flexibility in their ability to invade new cell types, tissues, and host species.

Unlike most other class I viral fusion proteins there is no crystal structure of intact S, severely limiting our mechanistic understanding of coronavirus fusion. While the basic structure of the S of SARS and SARS-2 can be revealed using cryo-EM techniques^6–8^ and part of the SARS-2 S2 subunit has been solved by X-ray crystallography^9,10^, many structural and functional aspects remain undetermined, especially since the active form of FPs (i.e. associated with membranes) is generally not determined in the crystallographic structure as they are hydrophobic and intrinsically disordered. ESR is a useful technique to study the effect of FPs on the membrane structure with implications for the mechanism leading to membrane fusion. It can also be used to determine the peptide structure in the membrane in the form of Pulse-Dipolar ESR^11,12^ and Power Saturation ESR^13^.

Previously, we used phospholipid (PC) spin labels to detect the perturbation of the membrane by viral FP and TMD^14–20^. We have shown that a wide range of viral FPs induce local membrane ordering, while their inactive mutants do not. We have interpreted this as associated with dehydration induced by the insertion of active FP^15,17,21^. We also found that this membrane ordering effect can be used as a criterion to identify the FP, as in the case of the gametic fusogen HAP2^20^. It was also used to identify the *bona fide* SARS-1 FP as the region immediately following the S2’ cleavage site, since the two other candidates do not induce local membrane ordering as efficiently. In the same study, we showed that the SARS-1 FP consists of FP1 and FP2 domains and that the activity of SARS-1 FP is Ca^2+^ dependent, with its FP1 and FP2 each binding one Ca^2+^ ion.^18^ Based on that finding, we proposed a bipartite model for the SARS-1 FP (fig 1A). We showed that the MERS-CoV FP also has FP1 and FP2 domains and its activity is also Ca^2+^ dependent, but only FP1 binds one Ca^2+^ ion^22^. This Ca^2+^ dependent fashion is also found in the Ebolavirus (EBOV) FP case. ^19^ In this study we apply the same methodology on the SARS-2 FP and compare it with the SARS-1 and MERS FP.

**Fig 1.**
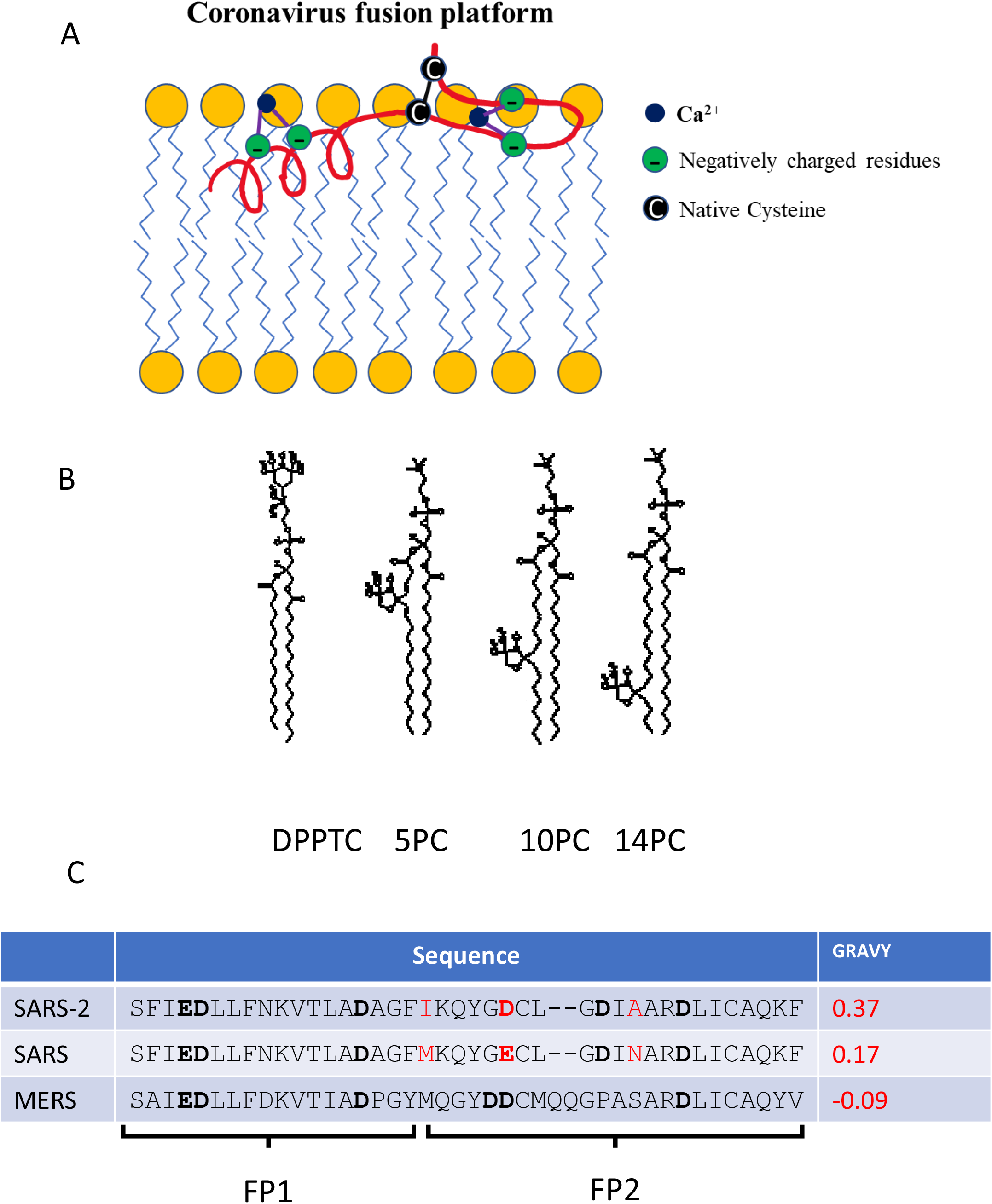
(A). A model of the coronavirus FP or “platform” interacting with a lipid bilayer (adopted from our previous model^18^ using the structural information from CD spectroscopy in this study, and the molecular dynamics model^26^). (B) The structure of spin labeled lipids used: DPPTC, 5PC, 10PC and 14PC. (C) The sequence alignment of SARS-2, SARS, and MERS FPs, and their grand average of hydropathy (GRAVY) values (calculated using GRAVY Calculator, http://www.gravy-calculator.de/index.php). The residues in red highlight the difference between SARS-2 and SARS FP. The residues in bold are the negatively charged residues which are potential Ca^2+^ binding sites.

## RESULTS

### Peptide Cloning and Expression

We previously identified the *bona fide* SARS-1 FP^23^ as the sequence that immediately downstream of the S2’ cleavage site. Based on the sequence alignment, we hypothesize that the homologous sequence on the SARS-2 S protein is the *bona fide* SARS-2 FP as well. We planned to test this by examining its membrane ordering effect as the criterion as before. We previously cloned the SARS-1 FP gene into a pET41 expression vector. In order to speed our research, we decided to generate the SARS-2 FP from this SARS-1 FP clone. As showed in Figure 1C, three residues are different between the SARS-1 FP and SARS-2 FP. Therefore, we have designed the primers and performed point mutagenesis on these three sites to generates the SARS-2 FP clone. The peptide was expressed in E. *Coli* and purified. Fig. 2A shows the SDS-PAGE gel of the fractions collected from His-tag column. The size of the peptide is correct. The combined fraction was further purified using size-exclusion chromatography. The size of the peptide was further validated by mass spectroscopy. All the data presented in this study are from this expressed version. We received synthesized SARS-2 FP from Biomatik USA LLC (Wilmington, DE) at a later time and have repeated some key experiments (CD, ESR and ITC) and found that the results between the expressed and synthesized ones are nearly identical. We used POPC/POPG/Chol=3/1/1 as the model membrane system. POPG is an anionic lipid and the POPC/POPG mixture is widely used in other fusion peptide research.^24,25^

**Fig 2.**
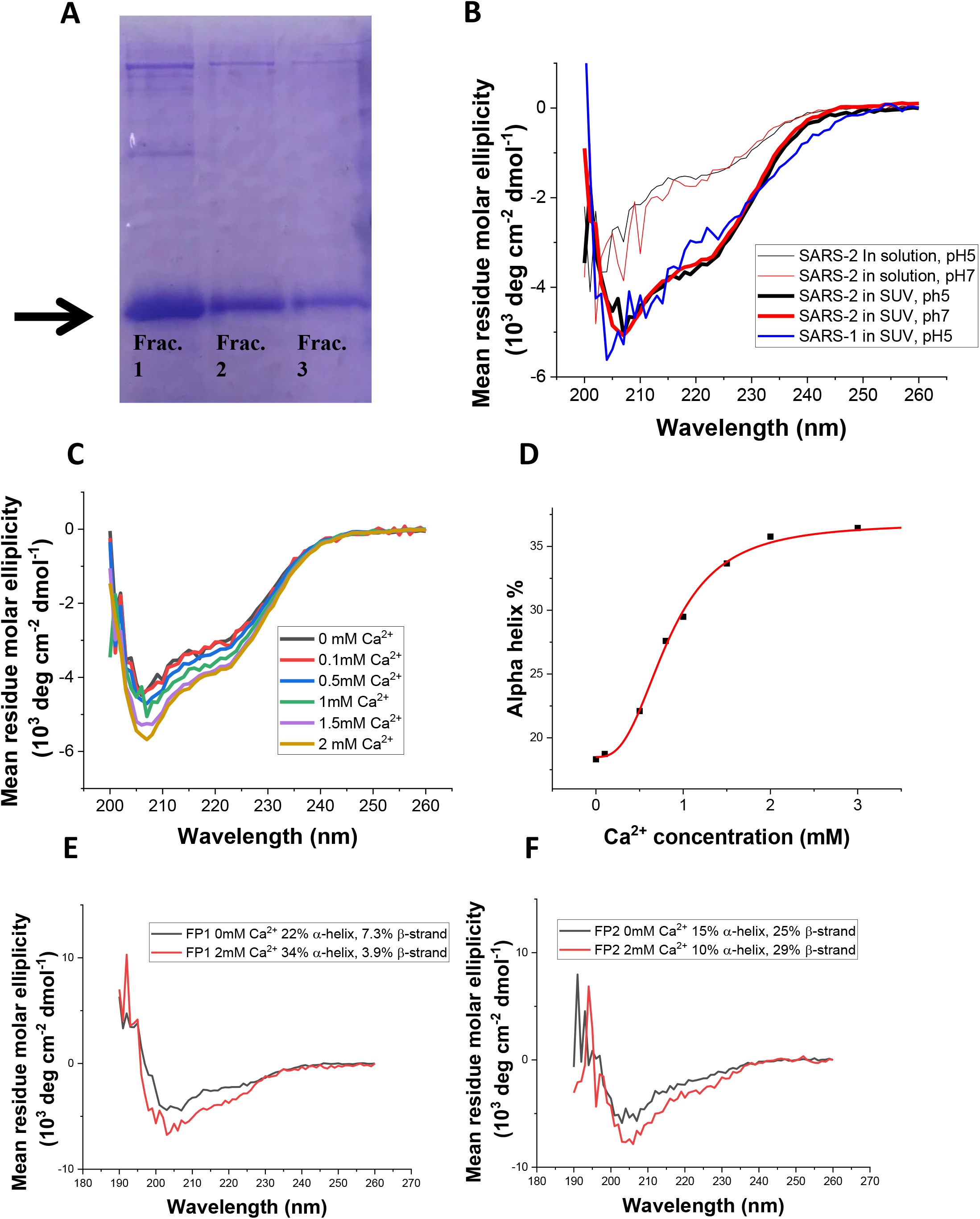
(A)The SDS-PAGE gel of the SARS-2 FP after purification using a His-tag affinity column; the arrow shows the bands for the FP. The fractions were the combined and further purified using size-exclusive chromatography. (B) CD Spectra of SARS-2 FP in solution (thin lines) and POPC/POPG/Chol=3/1/1 SUVs (thick lines) in pH 5 buffer (black) or pH 7 buffer (red) at 25°C, and SARS-1 FP in SUVs at pH5 (blue). (C) CD spectra for SARS-2 FP in SUVs at pH5 show that the alpha helix content increases with the increase of Ca^2+^ concentration. (D) The alpha helix content of the FP in membranes at different Ca^2+^ concentrations, which was calculated from (C) using K3D2 server^51^. The data were fit with the built-in logistic function of Origin, which shows the X_50_= 0.83 mM. (E, F) The CD spectra of SARS-2 FP1 (E) and FP2 (F) in SUV in pH5 buffer with 0mM (black) and 2 mM (red) CaCl_2_. The percentage of alpha helix and beta strand was calculated using K3D2.

### Circular dichroism spectroscopy (CD)

To determine whether the SARS-2 FP interacts with membrane, circular dichroism (CD) spectroscopy was used to determine the secondary structures of the peptides in solution and in membranes, which allows us to compare the structural transitions occurring between these two environments. As shown in fig. 2B, whereas the FP exhibits mostly a random coil structure in solution, it adopts a mixture of alpha-helix, beta-sheet and random-coil secondary structures in the presence of small unilamellar vesicles (SUVs) composed of POPC/POPG/Chol=3/1/1 and 1mM CaCl_2_ at pH 5. Here, we used SUVs instead of multilamellar lipid vesicles (MLVs) or large unilamellar vesicles (LUV) because MLVs and LUVs are too large for CD spectroscopy, since they would cause scattering and a noisy signal. We measured the CD spectra for SARS-1 FP under the same conditions, and found them similar to those of the SARS-2 FP, though the helical content of the SARS-2 FP is slightly greater in the membranes (27.6% for SARS-1, and 29.5% for SARS-2). To test the effect of pH on the secondary structure, we measured the CD spectra at pH5 and pH7. As shown in Fig 2B, the pH has little effect on the secondary structure of the SARS-2 FP in both the presence and absence of SUVs with 1 mM Ca^2+^. The structure in membranes consists of 29% alpha helix and 7% beta strand. To test the effect of Ca^2+^ on the secondary structure, we increased the Ca^2+^ concentration from 0 to 3 mM, and we found that the alpha helical contents of the SARS-2 FP increases from 18% to 35%, saturating around 2mM (cf. Fig 2C and D), which are for SUV’s and pH5. To identify the source of helical increase, we synthesized separately the FP1 domain and the FP2 domain and repeated the CD experiments. As shown in Fig 2E and 2F, FP1 has a higher percentage of alpha helical structure than beta strand, while FP2 has the opposite. This result is consistent with the newly published structural model of SARS-2 FP using molecular dynamics^26^. The presence of Ca^2+^ promotes the formation of alpha helix in FP1 but promotes the formation of beta strand in FP2. Thus, the CD spectra show that the SARS-2 FP is affected by binding to SUV membranes and Ca^2+^ has an overall effect of aiding the development of its secondary structure.

### SARS-2 FP Increases Membrane Ordering

The ESR signal of the spin labels attached to the lipids in membrane bilayers is sensitive to the local environment. Four spin labels were used: DPPTC has a tempo-choline headgroup and the spin is sensitive to changes of environment at the headgroup region; 5PC, 10PC and 14PC have a doxyl group in the C5, C10 or C14 position respectively of the acyl chain (Fig. 1B), and they are sensitive to the changes of local environment in the hydrophobic acyl chain region at the different depths. Using the NLSL software based on the MOMD model ^27,28^, the order parameter of the spin can be extracted, which is a direct measure of the local ordering of the membrane. Thus, the effect of peptide binding on the structure of the membrane can be monitored. These four spin-labeled lipids have been used in previous studies, and their ability to detect changes in membrane structure has been validated ^21,29^. Our previous studies examined the effect of various viral FPs, including those of influenza virus ^14,17^, HIV ^16^, Dengue Virus ^20^, Ebolavirus(EBOV)^19^, SARS-1^18^, and MERS^22^, as well as the FP of the ancestral eukaryotic gamete fusion protein HAP2 ^20^. All of these peptides were found to induce membrane ordering in the headgroup region as well as in the shallow hydrophobic region of bilayers (i.e., 5PC).

We used multilamellar vesicles (MLVs) in our ESR study as they can be much more concentrated than small unilamellar vesicles (SUVs) which greatly enhances the ESR signal. Furthermore, the use of MLVs is consistent with our previous studies ^16,17,20^. As shown in fig. 3A, for the MLVs composed of POPC/POPG/Chol=3/1/1, when the peptide:lipid ratio (P/L ratio) of FP increases from 0 to 5 mol%, the order parameter S_0_ of DPPTC increases significantly from 0.44 to 0.49 at pH 5 and in the presence of 1mM Ca^2+^ (with the effect of varying the Ca^2+^ concentration described in the next subsection). The increase of the S_0_ is similar to the effect of influenza FP as we have previously shown ^14–18,20,22^. The membrane order of 5PC also significantly increases from 0.53 to 0.65 at pH 5 when the P/L ratio increases (fig. 3B). The perturbation of the FP to the lipid bilayers exists down to 10 PC, although the effect is not as large (from 0.32 to 0.35, fig 3C). There is virtually no effect on the S_0_ of 14PC (cf, fig. 3D). The pH seems not to have a substantial effect from a comparison of results at pH5 and pH7, however acidification does boost the effect modestly (fig 3E). To show that the ordering effect is sequence-specific, we synthesized a peptide that has the same residues as the SARS-2 FP but in a shuffled order as a control. As shown in fig 3E, the shuffled-sequence peptide does not affect the membrane order at all. Clearly, the sequence of the FP is very important for its membrane ordering activity.

**Fig 3.**
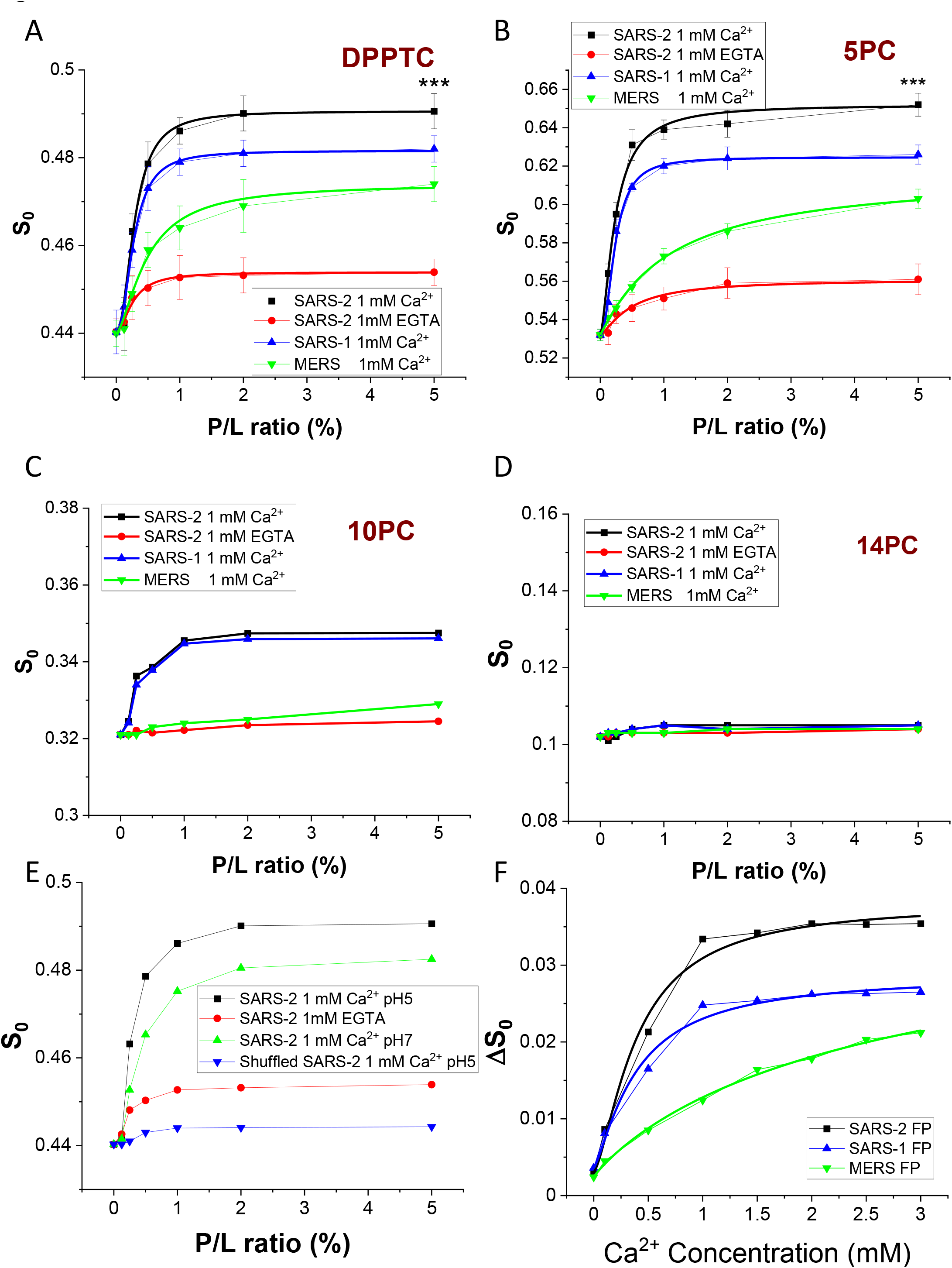
Plots of order parameters of DPPTC (A), 5PC (B), 10PC (C) and 14PC(D) versus peptide:lipid ratio (P/L ratio) of SARS-2 FP, SARS FP and MERS FP in POPC/POPG/Chol=3/1/1 MLVs in buffer at pH5 with 150 mM NaCl at 25°C. Black, SARS-2 FP with 1 mM Ca^2+^ and at pH 5; red, SARS-2 FP with 1 mM EGTA; blue, SARS FP with 1 mM Ca^2+^; green, MERS FP with 1 mM Ca^2+^. The curves were fitted with the logistic function (Table 1A and 1B) (E). Plots of local order parameters of DPPTC versus P/L ratio as in A-D with pH indicated. Black, SARS-2 FP at pH 5; red, SARS-2 FP triple mutant (E4A_D5A_D15A) at pH 5; blue, SARS-2 FP at pH 7; and green, a peptide with shuffled sequence of SARS-2 FP at pH 5. (F) Plot of local order parameters of DPPTC with and without 1% peptide binding (ΔS_0_) versus Ca^2+^ as in A-D. Black, SARS-2 FP; blue, SARS FP and green, MERS FP. The experiments were typically repeated three times. The typical uncertainties from simulation found for S_0_ range from 1-5 × 10^−3^, while the uncertainties from repeated experiments were 5-8 × 10^−3^ or less than ±0.01. We show in Panels A and B the bars for the standard deviation. Statistical significance analyses were performed using two-tailed Student’s *t*-test on the S_0_’s of 0% SARS-2 FP and 5% SARS-2 FP at the “1mM Ca^2+^” condition, *** ≤ 0.001, highly significant.

We also compared the effect of the SARS-2 FP to the SARS-1 FP and the MERS FP. As shown in figs. 3A and 3B, the shape of the S-curves of the SARS-2 and the SARS-1 FPs are similar in that they both saturate at around 2% P/L ratio, while the shape of the curve of the MERS FP is quite different in that it does not saturated even at 5% P/L ratio. It is worth mentioning that the 5% P/L ratio is very large in a natural scenario and that is why we did not increase the concentration further in our experiments. In an effort to quantify our results, we fit the S_0_ vs P/L ratio curves for DPPTC (fig 3A) and 5PC (fig 3B) using the logistic function 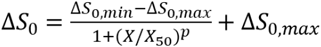 with fixed S_0,min_ and flexible S_0,max_, and also extracted the X_50_ and p from the fittings. The X_50_represents the P/L ratio for the FP to reach its half maximum effect, and the p value might be viewed as the cooperativity of the peptide when it induces membrane ordering. We compare the Max ΔS_0_, X_50_ and p values for the three CoV FP (Tables 1A and 1B, for DPPTC and 5PC respectively). We find that the maximal effect of the SARS-2 FP is greater than that of the SARS-FP (0.051 vs 0.039 for DPPTC and 0.12 vs 0.09 for 5PC) which is again greater than that of the MERS FP. The X_50_ values of the SARS-2 and SARS-1 FPs are almost the same (0.26 and 0.29 for DPPTC and 0.23 and 0.22 for 5PC), and they are both significantly smaller than that of the MERS FP (0.60 and 1.14 for DPPTC and 5PC, respectively). The p values for the SARS-2 FP are smaller than those for the SARS-1 FP but greater than those for the MERS FP. In conclusion, the SARS-2 fusion peptide shows a greater membrane ordering effect than does the SARS-1 FP and the MERS FP.

### The SARS-2 FP-Induced Membrane Ordering Is Calcium Dependent

We have previously found that the EBOV^19^, SARS-1^18^ and MERS^22^ FPs perturb the membrane structures in a Ca^2+^ dependent fashion. Because of the similarity between the SARS-1 and SARS-2 FPs, we expected to observe this Ca^2+^ dependency in the SARS-2 FP as well. For comparison we prepared a “calcium-free” buffer by not adding Ca^2+^ and instead adding 1 mM of the chelating agent EGTA to further remove traces of Ca^2+^. As shown in Figs 3A and B, the membrane ordering effects of the SARS-2 FP is suppressed. The small increase of S_0_ when the P/L ratio increases, we call “basal activity” for the SARS-2 FP, but this is insignificant compared to that in the presence of Ca^2+^.

In order to further examine the effects of Ca^2+^ on FP-inducing membrane ordering, we maintained the P/L ratio of the FP at a constant 1 mol%, and measured the order parameter S_0_ with increasing calcium concentrations ranging from 0 to 3.0 mM (fig. 3F); the highest calcium concentration used here is much higher than the extracellular concentration of Ca^2+^ in human adult lungs (ca. 1.3 mM) ^30^. The increase in S_0_ affected by the presence of both the FPs and the Ca^2+^; so, we generated a ΔS_0_-Ca^2+^ concentration plot, where ΔS_0_= S_0_ (membrane with 1% FP) − S_0_ (membrane without FP) vs. Ca^2+^ concentration. This subtraction cancels any membrane ordering induced just by Ca^2+^, with the ΔS_0_ at each Ca^2+^ concentration representing only the contribution of Ca^2+^ interacting with the FPs.

As shown in fig. 3F, Ca^2+^ increases the ΔS_0_ of all three CoV FPs. However, while the response pattern to the Ca^2+^ concentration is again similar within the SARS FPs, it is different from that of the MERS FP. We fit the ΔS_0_ vs Ca^2+^ concentration curves with the Hill equation 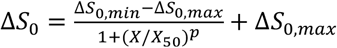 with fixed ΔS_0,min_ and flexible ΔS_0,max_ (Table 2), where X_50_ is the Ca^2+^ concentration that reaches 50% effectiveness, and the p value is the Hill slope interpreted as the cooperativity of the Ca^2+^ binding to the FP. While the X_50_ of the SARS-2 and SARS-1 FP are close (0.42 mM and 0.39 mM, respectively), the X_50_ of the MERS FP is much larger (2.85 mM). The Hill slopes for SARS-2 and SARS FP are 1.67 and 1.30 respectively, indicating a positive cooperative binding of Ca^2+^ to the FP, while the Hill slope for the MERS-FP (0.90) is appears not to be cooperative. We have shown previously that the SARS-1 FP binds two Ca^2+^ ions, each on FP1 and FP2^18^, and the MERS FP binds only one Ca^2+^ ion, possibly on FP1^22^. The difference between the Hill slopes appears to reflect this difference.

### Enthalpy Changes of FP-Membrane Interactions

The importance of Ca^2+^ can be further studied by comparing membrane binding enthalpies of FPs as measured by isothermal titration calorimetry (ITC) in the presence and absence of Ca^2+^. In this experiment, small amounts of peptide were injected into a reaction cell containing a large excess of SUVs. Thus, during the whole titration process the amount of available membrane can be regarded as constant, and all injected peptides can be regarded as binding to the membranes. As a result, the reaction heat in each injection is equal. The enthalpy of reaction can be calculated from the average of heat in each injection^31,32^. We performed this experiment in the buffer with or without 1 mM Ca^2+^. We then compared the difference in reaction enthalpy, which is caused by the presence of Ca^2+^.

As shown in Table 3A, the binding enthalpy of the CoV-FPs is greater in the presence of Ca^2+^ than in its absence. In both cases, the ΔH for SARS-2 and SARS-1 FP is greater than the MERS FP for this exothermic reaction with that for SARS-2 being significantly greater than for SARS-1 only in the presence of Ca^2+^. When the difference between the with/without Ca^2+^ cases was compared (ΔΔH ≡ ΔH (in presence of Ca^2+^) – ΔH (in absence of Ca^2+^)), the ΔΔH of SARS-2 is greater than that of SARS-1 FP, which is greater than that for MERS FP. The enthalpy of FP-membrane binding mainly results from the folding of the FP^33,34^. Thus, the results suggest that: 1) SARS-2 FP folding is stronger than that of the SARS-1 FP and even more so than that of the MERS FP; 2) While Ca^2+^ appears to promote the folding of the FP, it does it more effectively for the two SARS FPs than the MERS FP. That Ca^2+^ promotes SARS-2 FP folding is consistent with the CD results (Fig 2C and 2D) which show the alpha helical content increases as the [Ca^2+^] increases reaching a plateau for [Ca^2+^]>2mM. This may be compared with the ESR observation that the Ca^2+^ promotes FP-induced membrane ordering in a similar [Ca^2+^] dependent manner (compare Fig 3A,B with Fig 2D).

### Interactions of FPs with Calcium Ions Detected by ITC

We also used ITC to investigate whether Ca^2+^ ions directly interact with SARS-2 FPs. During this titration, the Ca^2+^ was titrated into the reaction cell with FPs in solution. Substantial heat absorbed during the titration was observed, with the heat absorbed saturating toward the end of the titration. As shown in fig. 4 and Table 3B, when the ΔH vs. Ca^2+^:FP molar ratio plot was fitted using a one-site model, that makes the simple assumption that all binding sites have the same binding affinity, the enthalpy change is ΔH=7.2 kCal/mol, with a binding constant K_b_=2.7×10^4^ M^-1^ and stoichiometry n=1.86. From these parameters, we further calculated the free energy change ΔG=− RTlnK_b_=6.3 kCal/mol and − TΔS= ΔG− ΔH = − 0.9 kCal/mol. Thus, the calcium-SARS-2 FP interaction is endothermic, and the binding ratio is two calcium ions per peptide. The reason why the reaction is endothermic could be because the buffer solution (150 mM NaCl) contains large amounts of Na^+^ ions that could also interact with FP and that a stronger binding of Ca^2+^ cations displaces bound Na^+^ into the bulk solution. To test this hypothesis, we reduced the NaCl concentration to 50 mM and performed an ITC titration. However, the baselines of the titration curve we obtained were unstable, and we could not extract meaningful data. The reason for this result is unclear, but it is likely that the peptide requires a certain level of salt concentration to maintain its structure.

**Fig 4.**
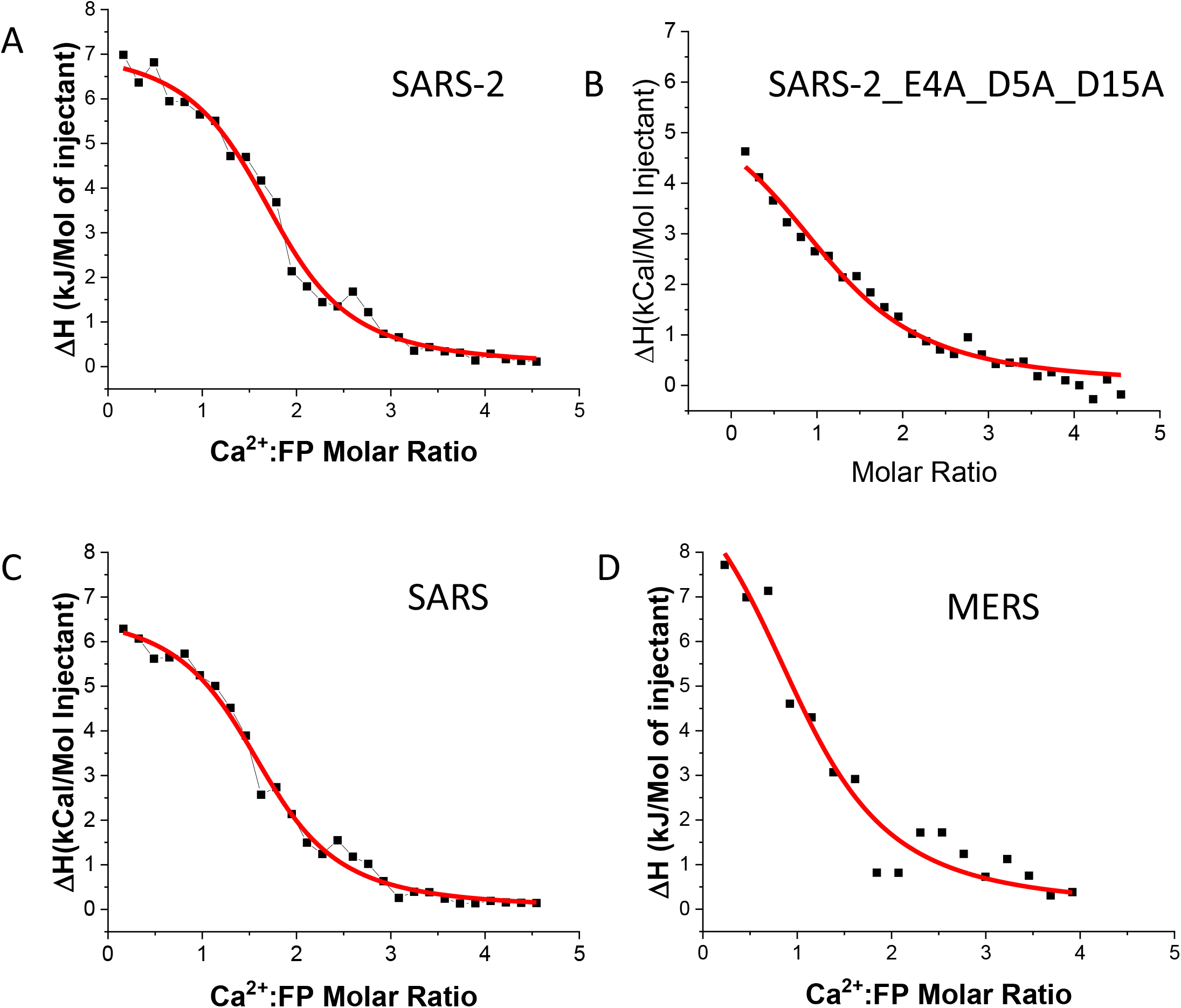
ITC analysis of Ca^2+^ binding to FP of the coronaviruses. (A) SARS-2 FP, (B) SARS-2 FP triple mutant (E4A_D5A_D15A). (C) SARS-1 FP. (D) MERS FP. The peptides were titrated with CaCl_2_ in the pH5 buffer. The integrated data represent the enthalpy change per mole of injectant, ΔH, in units of kcal/mol as a function of the molar ratio. The data were fitted using a one-site model (parameters shown in table 3B). Data points and fitted data are overlaid.

We performed the same experiment with SARS-1 FP and MERS FP (fig. 4C and 4D). As shown in Table 3B, like SARS-2 FP, SARS-1 FP also exhibits an endothermic reaction with Ca^2+^, with a stoichiometry of n=1.70, ΔH=6.8 kCal/mol, and K_b_=2.5×10^4^ M^-1^. For the MERS FP, the stoichiometry is n=1.12, suggesting that it only binds one Ca^2+^, and the ΔH (3.1 kCal/mol) is significantly lower than that of the SARS-1 FPs, also resulting from the less Ca^2+^ binding. We also generated a triple mutation on the SARS-2 FP (E4A_D5A_D15A), which completely removed the potential Ca^2+^ binding site on the FP1 section. As shown in fig. 4B and Table 4B, the n is reduced to 1.03, which is coincident with our hypothesis that the triple mutant can only bind one Ca^2+^ ion in its FP2 section. In conclusion, these ITC experiments demonstrate strong evidence for direct calcium-FP interactions, and we associate it with one Ca^2+^ cation binding per FP1 or FP2.

### The Ca^2+^ dependency is highly specific

We have already shown that the SARS-2 FP induces membrane ordering in a Ca^2+^ dependent fashion. But how specific is it? In order to address this question, we have repeated the ESR experiment in the presence of a number of different ions, including most of the biologically relevant ions (Mg^2+^, Al^3+^, K^+^, Fe^2+^, Fe^3+^, Zn^2+^, Mn^2+^) as well as Tb^3+^ which is not biologically relevant but is widely used as a Ca^2+^ substitute as the size is similar to the Ca^2+ 35^. We used the 1mM EGTA case as a control; as mentioned above even without any Ca^2+^, the FP has a “basal” activity. We did not measure the effect of Na^+^ as there is 150 mM Na^+^ in the buffer, which provides the necessary ionic strength for the folding of the FP. As shown in Fig 5A, the membrane ordering activity of the FP in the presence of all other ions except Tb^3+^ is significantly lower than that in the presence of Ca^2+^. The Tb^3+^ case is the closest to the Ca^2+^ case, but still substantially smaller. The K^+^ has almost zero effect on the activity; it is almost the same as the EGTA case. When the concentration of some ions increases to 2mM (fig 5B), the trends are the same. We then fixed the P/L ratio to 1% and increased the concentration of ions (fig 5C), we again find that the Ca^2+^ and Tb^3+^ will allow the FP to function more effectively (as measured by ΔS_0_) in higher concentrations, while the Mg^2+^ and Mn^2+^ only boost the effect slightly. We repeated the ion-FP ITC experiments for several ions, and one sees from fig 5D, that while the Tb^3+^ curve is comparable to that of the Ca^2+^, the other ions have a significantly lower reaction heat than the Ca^2+^, which again shows the specificity of Ca^2+^-FP binding.

**Fig 5.**
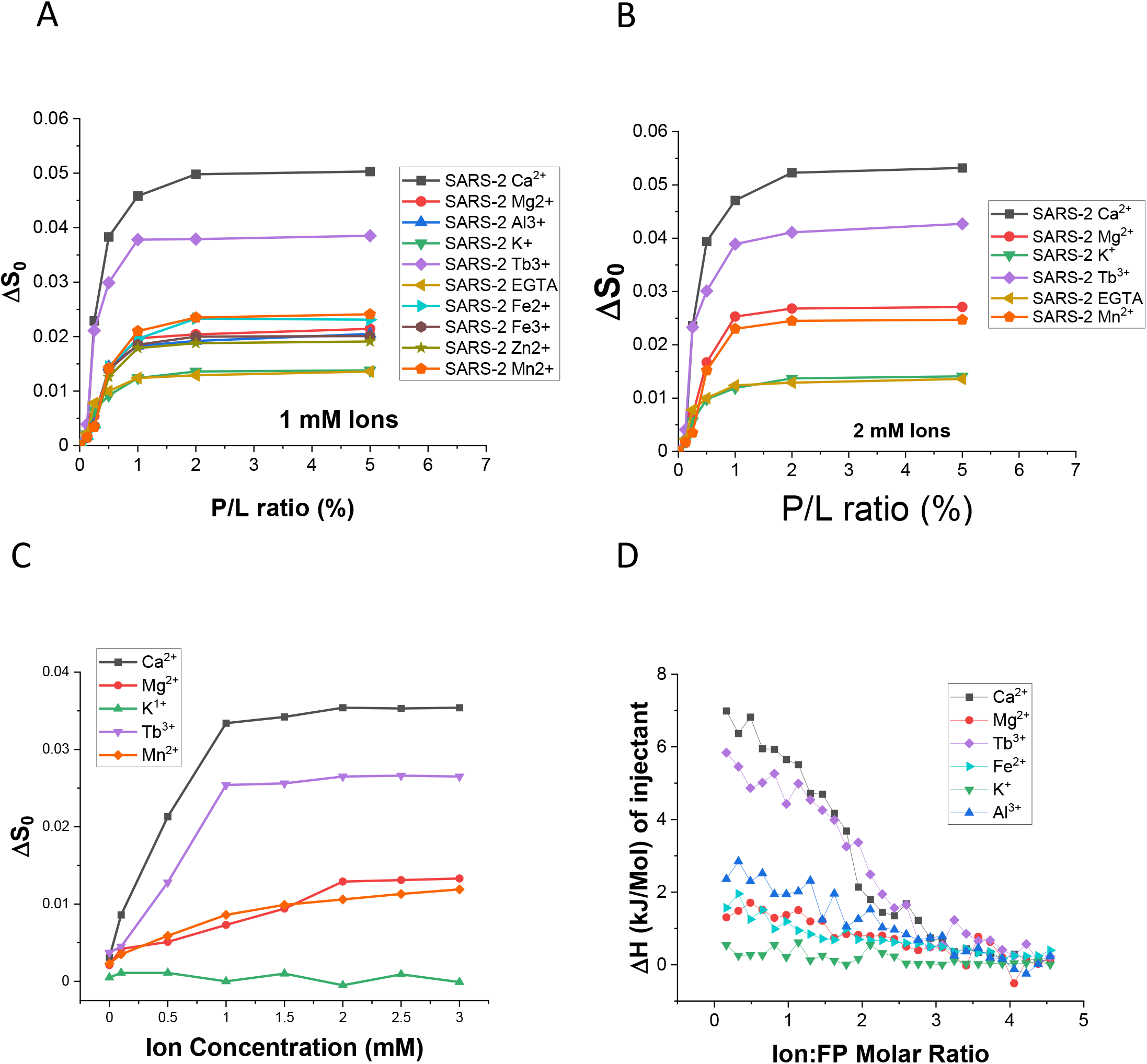
The specificity of Ca^2+^. (A) Plots of local order parameter changes of DPPTC (ΔS_0_) versus the P/L ratio with 1 mM ion concentration for a variety of ions and in buffer with 150 mM NaCl at pH5 and at 25°C. (B) The same with 2 mM ion concentration. (C) Plots of local order parameter changes of DPPTC (ΔS_0_) versus the concentration of the ions with 1% P/L ratio and at pH5. (D) ITC analysis of ions binding to SARS-2 FP in solution. The peptides were titrated with CaCl_2_ in pH5 buffer.

## DISCUSSION

The mechanism of membrane fusion is still only partially understood. Our past work has shown that numerous viral FPs induce increased membrane ordering in a collective fashion, i.e. a significant increase in S_0_ as a function of the P/L ratio^14,15,17–20,22,36^. We further suggested that FP-induced membrane ordering is accompanied with dehydration resulting from peptide insertion, and this is a prerequisite step for removal of the repulsive forces between two opposing membranes, thereby facilitating initialization of membrane fusion ^14–17,20^. We, in conjunction with our collaborators, have also determined that the CoV (SARS-CoV and MERS CoV) and EBOV requires Ca^2+^ for viral entry and that the Ca^2+^ binding site is on the FP^18,19^. Our current study extends these observations to the SARS-2 FP revealing several new features.

We have established the hypothesis^23^ that the sequence immediately downstream from the S2’ cleavage site is the *bona fide* SARS-1 FP ^18^. Thus, we predicted that the homologous sequence on the SARS-2 S protein to the SARS-1 FP is also the *bona fide* SARS-2 FP. We have now shown that this SARS-2 FP does effectively induce membrane ordering, thereby confirming it must be the FP for the SARS-2 S protein. We also showed that it perturbs the membrane structure in a Ca^2+^ dependent fashion as do the SARS-1 and MERS FPs.

Although this result was expected considering the similarity between SARS-2 and SARS-1 FPs, we did find that SARS-2 FP leads to greater membrane ordering than SARS-1 FP, especially in the head group and upper hydrophobic regions. It both induces a greater degree of membrane ordering and has smaller X_50_ values. The SARS-2 FP also has a greater level of response to the Ca^2+^ concentration (cf. fig 3F and Table 2) as it has a greater Max(ΔS_0_) and the Hill slope is greater. This is supported by the fact that that the SARS-2 FP has a higher binding enthalpy than the SARS-1 FP (cf. Table 3A). All the above suggest that the SARS-2 FP is more effective in membrane binding and Ca^2+^ binding than SARS-1 FP.

Since there is only a three residue difference between the SARS-2 and SARS-1 FP, the most reasonable cause for the difference between the activity of these two FPs must be in these three residues. We suspect that the greater hydrophobicity of the three residues in SARS-2 FP is the cause. The hydropathy index for Ile (4.5), Asp^-^ (−3.5) and Ala (1.8) of the SARS-2 FP is greater than or equal their counterparts (Met (1.9), Glu^-^ (−3.5) and Gln (−3.5)) of the SARS-1 FP (greater positive value means more hydrophobic) ^37^. Thus, the grand average of hydropathy value (GRAVY) for SARS-2 FP is also greater than the SARS-1 FP (cf. Fig 1C). In the case of peptide-membrane interaction, the interface scale of hydrophobicity is more useful as it measures the free energies of transfer ΔG (kCal/mol) from water to POPC interface^38^. The interface scale of the Ile (−0.31), Asp^-^ (1.23) and Ala (0.17) of the SARS-2 FP is also greater than their counterparts (Met (−0.23), Glu^-^ (2.02), and Gln (0.58)) of the SARS-1 FP^38^ (here more negative value means more hydrophobic), thus the overall interface scale value for the SARS-2 FP is also significantly more hydrophobic than the SARS-1 FP.

In the process of the FP interacting with the membrane, most of the driving force comes from the enthalpy gain by the folding of FP, whereas the opposing force comes from the entropy loss of restricting the FP on the membrane surface.^24,33,34^ The latter can be partly compensated by releasing water molecules during a hydrophobic interaction between the FP and the hydrophobic region of the membrane. The more hydrophobic SARS-2 FP interacts with the hydrophobic core more strongly, inserts more deeply into the membrane, and likely releases more water molecules in the headgroup region and thus reduces the entropy loss more effectively than does the SARS-1 FP. SARS-2 FP also has a greater effect on the shallower hydrophobic region (5PC, fig 3B). The folding of the FP is also modulated by how deep it inserts into the membrane. We observed that the enthalpy release of the SARS-2 FP-membrane insertion is greater, indicating better folding than the SARS-1 FP in the membrane, which also increases the driving force of the FP-membrane interaction.

These mutations also account for the better Ca^2+^-FP binding and the better Ca^2+^ response of the SARS-2 FP. Asp has been shown to have a greater Ca^2+^ binding affinity than Glu in aqueous solution.^39^ Although Ala may not directly be involved in the interaction with Ca^2+^ in this case, it has a greater Ca^2+^ affinity than Gln even at high pH.^40^

The greater ordering effect of the SARS-2 FP implies that the SARS-2 CoV enters the host cells more effectively than SARS-1-CoV if other factors remain the same. This may be one of the reasons why the SARS-2-CoV invades host cells more easily and cause more severe impact to public health. The greater [Ca^2+^] response also implies that control of its access to Ca^2+^ could reduce its infectivity.

We further investigated how specific is the role of Ca^2+^ in the activity of the SARS-2 FP. By comparing a series of ions, we found that the FP-Ca^2+^ interaction is very specific. Although most other ions can enhance the membrane ordering slightly above the “basal activity”, their effect is much smaller than that of the Ca^2+^. The ion-FP interaction is also weaker, as revealed by ITC. Although the FP has negatively charged residues and should attract cations, this selectivity is likely to be governed by the many-body polarization effect^41^, which is an energetic cost arising from the dense packing of multiple residues around the metal ion. The many-body effect depends on the size of the ion, as well as the number and spatial arrangement of the residues around it. There are three negatively charged residues in either the FP1 or FP2 region which satisfies the requirement for the requirement of a multi-body interaction. We also showed that the only close alternative candidate to Ca^2+^ is Tb^3+^, which has a size that is similar to that of Ca^2+^ (radius of Ca^2+^ and Tb^3+^ in solution is 103 and 101 picometer respectively). We found the ion-FP interaction is endothermal. We suspect that the substantial amount of Na^+^ already in solution initially interacts with the charged residues, and there is a more effective replacement by Ca^2+^ during the titration. That is, ions with suitable size can efficiently replace the Na^+^, but this suggestion requires verification.

Calcium ions are important modulators of membrane fusion. Because of their positive charges, they possess a generally enhancing but indirect effect on membrane fusion by electrostatic interactions with negatively charged headgroups of lipid bilayers, and thus decrease the electrostatic repulsion of the two opposing membranes that are in close proximity prior to undergoing fusion. Calcium ions can also directly interact with fusion protein machineries, thereby activating their fusogenicity, and in such cases membrane fusion is clearly calcium-dependent. For example, in cellular SNARE-mediated synaptic vesicle fusion, calcium ions have been shown to be required activators for such fusion machinery ^42–44^, i.e. without calcium ions present, membrane fusion does not take place. Thus, SNARE-mediated synaptic membrane fusion is considered to be calcium-dependent whereby calcium ions are fusion-activators. There are other situations in which calcium ions have been shown to interact with protein fusogens and enhance their fusogenicity; but without calcium, membrane fusion can still occur albeit at a reduced rate. In such cases membrane fusion is only partly dependent on calcium ions.

There are relatively few cases for which calcium plays a direct role in viral membrane fusion, especially for an N-terminal FP. In the case of influenza and HIV with an N-terminal FP, calcium plays only an indirect enhancing effect on membrane fusion by affecting the membrane only^45^. However, in cases such as CoV, calcium also plays a direct role by interacting with the FP directly. The infection of Rubella virus has been shown to require Ca^2+^, but its E1 envelope glycoprotein is a class II viral fusion protein that has an internal FP^46^. We have shown that the EBOV GP protein is another case for Ca^2+^ dependence, but EBOV FP is an internal one^19^. In this study and our previous study^18,22^, we showed that the viral entry of members of the CoV virus family (SARS-1, SARS-2 and MERS), which all have an N-terminal FP, are also Ca^2+^ dependent, Ca^2+^ has a direct activating role, and the FP is the Ca^2+^ target. Insights into the Ca^2+^ dependency of the FP would help to better understand the mechanism of the SARS-2 infection, and should also help to determine whether the membrane fusion of SARS-2 occurs on the cell surface or inside the endosome by controlling the exocellular or intracellular [Ca^2+^]. This could lead to therapeutic solutions to either targeting the FP-calcium interaction, or block the Ca^2+^ channel, and also repurpose already approved drugs for the ongoing COVID19 pandemic.

## MATERIALS AND METHODS

### Lipids and Peptides

The lipids POPC, POPG, and the chain spin labels 5PC, 10PC and 14PC and the head group spin label dipalmitoylphospatidyl-tempo-choline (DPPTC) were purchased from Avanti Polar Lipids (Alabaster, AL) or synthesized in our laboratory according to previous protocols. Cholesterol was purchased from Sigma (St. Louis, MO). The shuffled sequence of SARS-2 was generated using Sequence Manipulation Suite (http://www.bioinformatics.org/sms2/shuffle_protein.html), and this peptide was synthesized by Biomatik. The sequences of the peptides and the structure of the spin labeled lipids are shown in fig. 1C.

We previously cloned the SARS-1 FP gene into a pET31 expression vector and the SARS-2 FP gene is generated by the point-mutagenesis from the SARS-1 FP clone. The mutations were generated using a USB Change-IT site directed mutagenesis kit (Affymetrix). The sequence has been confirmed in the sequencing service provided by the Biotechnology Core Facility (BRC) at Cornell University. The protocol of expression and purification also follows published procedures^19,47^. Briefly, the relevant plasmids were transformed in BL21(DE3) Escherichia coli competent cells and grown at 37 °C to an optical density ∼0.8. Protein expression was induced for 3 hours at 30°C by 0.5 mM IPTG. The harvested cells were lysed by sonication and clarified by centrifugation at 40,000 rpm for 45 minutes. The supernatant containing the His-tagged fusion protein was transferred to a pre-equilibrated Ni-NTA agarose resin column, and the supernatant and resin were incubated for 2 hours at 4 °C on a rotator in wash buffer (containing 25 mM Tris, 500 mM NaCl, 20 mM Imidazole, 5% glycerol, 5 mM β-ME, and 10 mM CHAPS, pH 8). The resin was then rinsed with digestion buffer (containing 25 mM Tris, 50 mM NaCl, 5 mM CaCl2, and 5% glycerol, pH 7.5). 125 µL of Factor Xa (1 mg/mL) in 15 mL digestion buffer was added to the resin and incubated overnight at room temperature. The proteins were eluted using 50 mL wash buffer, dialyzed against dialysis buffer (25 mM Tris, 50 mM NaCl, 5% Glycerol, pH 8.5), and purified using a Superdex Peptide 10/300 GL gel-filtration column (GE Healthcare).

### Vesicle Preparation

The composition of membranes used in this study is consistent with our previous study ^23^. The desired amount of POPC, POPG, cholesterol and 0.5% (mol:mol) spin-labeled lipids in chloroform were mixed well and dried by N_2_ flow. The mixture was evacuated in a vacuum drier overnight to remove any trace of chloroform. To prepare MLVs, the lipids were resuspended and fully hydrated using 1 mL of pH 7 or pH 5 buffer (5 mM HEPES, 10 mM MES, 150 mM NaCl, and 0.1 mM EDTA, pH 7 or pH 5) at room temperature (RT) for 2 hours. To prepare SUVs for CD and ITC measurements and the PP-SUV system, the lipids were resuspended in pH 7 or pH 5 buffer and sonicated in an ice bath for 20 minutes or when the suspension became clear. The SUVs solution was then further clarified by ultracentrifugation at 13,000 rpm for 10 min.

### Circular Dichroism (CD) Spectroscopy

The CD experiments were carried out on an AVIV CD spectrometer Aviv Model 215. The peptides were mixed with SUVs in 1% P/L ratio with a final peptide concentration of 0.1 mg/mL at RT for more than 10 min before the measurements. The measurements were performed at 25°C and two repetitions were collected. Blanks were subtracted and the resulting spectra were analyzed. The mean residue weight ellipticity was calculated using the formula: [Θ] = θ/(10×c×l×n), where θ is the ellipticity observed (in degrees); c is the peptide concentration (in dmol); l is the path length (0.1 cm); and n is the number of amino acids per peptide ^32^.

### Isothermal Titration Calorimetry (ITC)

ITC experiments were performed in an N-ITC III calorimeter (TA Instrument, New Castle, DE). To measure the enthalpy of FP membrane binding, FP at 20 µM was injected into 1 mL 5 mM SUV solution at 37°C. Each addition was 10 µL, each injection time was 15 sec, and each interval time was 5 min. Each experiment comprised about 25 to 30 injections. The data were analyzed with Origin (OriginLab Corp., Northampton, MA).

To measure the peptide-Ca^2+^ interaction a total of 300 μL 2 mM CaCl_2_ in pH 5 buffer was injected into 0.4 mM FP in pH 5 buffer at 37°C in a stepwise manner consisting of 10 µL per injection except that the first injection was 2 μL. The injection time was 15 sec for each injection and the interval time was 10 min. The background caused by the dilution of CaCl_2_ was subtracted by using data from a control experiment that titrated CaCl_2_ in a pH 5 buffer. The data were analyzed with Origin. The one-site model was used in the fitting to calculate the thermodynamic parameters. The protein concentration is determined by dry weight and UV spectroscopy at wavelength = 280 nm with extinction coefficient ε=1520 M^-1^ cm^-1^ for SARS-2 and SARS-1 FP, and 4080 M^-1^ cm^-1^ for MERS FP in the oxidized condition.

### ESR spectroscopy and nonlinear least-squares fit of ESR spectra

To prepare the samples for lipid ESR study, the desired amounts of FPs (1 mg/mL) were added into the lipid MLVs dispersion. After 20 min of incubation, the dispersion was spun at 13,000 rpm for 10 min. The concentrations of peptide were measured using UV to ensure complete binding of peptide. The pellet was transferred to a quartz capillary tube for ESR measurement. ESR spectra were collected on an ELEXSYS ESR spectrometer (Bruker Instruments, Billerica, MA) at X-band (9.5 GHz) at 25°C using a N_2_ Temperature Controller (Bruker Instruments, Billerica, MA).

The ESR spectra from the labeled lipids were analyzed using the NLLS fitting program based on the stochastic Liouville equation ^28,48^ using the MOMD or Microscopic Order Macroscopic Disorder model as in previous studies ^14–17,20^. The fitting strategy is described below. We employed the Budil *et al*. NLLS fitting program ^28^ to obtain convergence to optimum parameters. The g-tensor and A-tensor parameters used in the simulations were determined from rigid limit spectra ^14^. In the simulation, we required a good fit with a small value of χ^2^ and also good agreement between the details of the final simulation and the experimental spectrum. Each experiment (and subsequent fit) was repeated 2 or 3 times to check reproducibility and estimate experimental uncertainty. Two sets of parameters that characterize the rotational diffusion of the nitroxide radical moiety in spin labels were generated. The first set is the rotational diffusion constants. R_⊥_and R_||_ are respectively the rates of rotation of the nitroxide moiety around a molecular axis perpendicular and parallel to the preferential orienting axis of the acyl chain. The second set consists of the ordering tensor parameters, S_0_ and S_2_, which are defined as follows: S_0_=<D_2,00_>=<1/2(3cos2θ− 1)>, and S_2_=<D_2,0_^2+^D_2,0-2_> =<√(3/2)sin2θcos2φ>, where D_2,00_, D_2,02_, and D_2,0-2_ are the Wigner rotation matrix elements and θ and φ are the polar and azimuthal angles for the orientation of the rotating axes of the nitroxide bonded to the lipid relative to the director of the bilayer, i.e. the preferential orientation of lipid molecules; the angular brackets imply ensemble averaging. S_0_ and its uncertainty were then calculated in standard fashion from its definition and the dimensionless ordering potentials C_20_ and C_22_ and their uncertainties found in the fitting. The typical uncertainties we find for S_0_ range from 1-5 × 10^−3^, while the uncertainties from repeated experiments are 5-8 × 10^−3^ or less than ±0.01. S_0_ indicates how strongly the chain segment to which the nitroxide is attached is aligned along the normal to the lipid bilayer, which is believed to be strongly correlated with hydration/dehydration of the lipid bilayers. As previously described, S_0_ is the main parameter for such studies ^17,49,50^.

## Acknowledgement

This work was funded by NIH grants R01GM123779 and P41GM103521. We thank Dr. Susan Daniel and Dr. Gary Whittaker for providing numerous helpful discussions and suggestions. We also thank Dr. Harel Weinstein and Dr. George Khelashvili for sharing the results of their molecular dynamics study.

## Abbreviations

CD: circular dichroism
CoV: coronavirus
ESR: electron spin resonance
FP: fusion peptide
ITC: isothermal calorimetry
LUV: large unilamellar vesicle
MERS: Middle East Respiratory Syndrome
MLV: multilamellar vesicle
MOMD: microscopic order but macroscopic disorder
POPC: 1-palmitoyl-2-oleoyl-glycero-3-phosphocholine
POPG: 1-palmitoyl-2-oleoyl-sn-glycero-3-phospho-(1’-rac-glycerol)
SARS: Severe Acute Respiratory Syndrome
SUV: small unilamellar vesicle
TMD: transmembrane domain

## Notes

### Competing Interest Statement

The authors have declared no competing interest.

